# Sugar metabolism changes in response to the ultraviolet B irradiation of peach (*Prunus persica* L.)

**DOI:** 10.1101/145870

**Authors:** Xuxu Wang, Xiling Fu, Min Chen, Lei Huan, Wenhai Liu, Yangang Gao, Wei Xiao, Xiude Chen, Ling Li, Dongsheng Gao

**Author notes:** These authors contributed equally to this work (Xuxu Wang, Xiling Fu). These authors parallel correspondence.

## Abstract

The protected cultivation of peach (*Prunus persica* L.) trees is more economical and efficient than traditional cultivation, resulting in increased farmers’ incomes, but the peach sugar contents are lower than in open planting. In the greenhouse, a high-sugar variety of peach ‘Lumi 1’ was irradiated with 1.44 KJ·m^−2.^d^−1^ intensity ultraviolet B radiation. The soluble sugar contents in fruit, peel and leaf were quantified using liquid chromatography. Overall, sucrose and sorbitol increased before the second fruit-expansion period. To further understand the mechanisms regulating sucrose and sorbitol accumulation in peach fruit, expression profiles of genes involved in sugar metabolism and transport were measured. The activity and translocation protein contents of these enzymes were measured by enzyme-linked immunosorbent assay. The increased sucrose synthase activity and sucrose transporter level in the pericarp promoted the synthesis of sucrose and intake of sucrose into fruit. Sorbitol transport into fruit was promoted by the increased sorbitol transporter protein levels in leaves. In summary, greenhouse the sucrose and sorbitol contents were increased when supplemented with 1.44 kJ·m^−2^·d^−1^ ultraviolet B radiation before the second fruit-expansion period of peach.

## Introduction

The peach (*Prunuspersica* L.) originated in China have been grown there for 3,000 years (Zheng et al., 2014). The protected cultivation of fruit trees is more economical and efficient than traditional cultivation, resulting in increased farmers’ incomes and the economic development of rural areas (Wang et al., 2016). Therefore, the protected cultivation of fruit trees in China has increased in recent years (Wang et al., 2016).

However, cultivated plants are restricted by light, ultraviolet (UV) radiation, ventilation and other environmental factors (Campbell et al., 2005). Thus, the protected cultivation of peach trees reduces the fruits’ flavor and soluble solid content compared with outdoor cultivation under natural conditions (Desnoues et al., 2014; Rolland et al., 2006). This seriously affected the profitability of the peach industry and its sustainable development, which do not meet consumer expectations (Crisosto, 2002). Therefore, it is important to improve the adaptability of peach to the greenhouse environment and improve the fruit quality.

Sweetness is an important component determine fruit quality which depends on the sugar contents and composition (Kroger et al., 2006). ‘Lumi 1’ (*Prunuspérsica* L. cv. Lumi 1) is a high-sugar variety with a soluble solids content of 13%. It was bred in our laboratory, and is a bud mutation of ‘Snow Kist’, a USA peach variety. The peel is dark red with white flesh, with desired hardness, adhesion, flavor and fragrance. Its vitamin C content was 6.98 mg·100 g^−1^. Thus, it has excellent qualities, as well as good storage and transportation performances.

The photosynthetic products of rosaceae plants, such as apple (Filip et al., 2016), cherry (Gao et al., 2003), loquat (Suzuki et al., 2014), peach (Bianco et al., 2000) and pear (Quro et al., 2000), are transported by the main metabolites of sorbitol and sucrose into the fruit (Yamaki, 2010). In higher plants, sugars are important structural materials and energy sources, and they also affect growth and development throughout the plant’s life cycle, playing central roles in germination, flowering and senescence (Loreti et al., 2001; Yao et al., 2011; Leskow et al., 2016). Sugars also have signaling functions (Dekkers et al., 2004; Kato-Noguchi et al., 2010; Rolland et al. 2006).

Important biochemical indices include the activity levels of important enzymes, which are strongly correlated to sucrose metabolism (Beruter and Studer Feusi, 1995). In peach or other Rosaceae plants, sucrose, sorbitol, glucose and fructose are mainly transported into the vacuoles by sucrose transporter (SUT), sorbitol transporter (SOT), tonoplastic monosaccharide transporter (TMT) and sugar transporter protein (STP), which are special transporter proteins located on the vacuole membrane (Hu et al., 2014; Sauer, 2007; Büttner, 2007). The SUT mainly mediates the transport of sucrose and maltose (Sauer, 2007), and the SOT mainly mediates the transport of sorbitol (Hu et al., 2014). The TMT and STP mainly mediate the transmembrane transport of glucose, fructose and lactose (Büttner, 2007). Sucrose enters the cell as sucrose or as hexose and is first hydrolyzed into glucose and fructose by cell wall invertase (CWINV) (Persia et al., 2008), which is related to plant stress resistance (Albacete et al., 2015). In sink cells, the transported sugars are either metabolized or stored. In the cytoplasm, there are a variety of enzymes involved in sugar metabolism that form a complex regulatory network, including neutral invertase (NINV), sucrose phosphate synthase (SPS) (Hashida et al., 2016; Yativ et al., 2010) and sucrose synthase (SUS) (Klotz and Haagenson, 2008). Vacuolar acid invertase (VAINV) can convert sucrose to fructose and glucose (Lin et al., 2015) after sucrose is transported into the vacuole (Chen et al., 2017). Sorbitol is synthesized in source tissues, such as chloroplast cells, by sorbitol-6-phosphate dehydrogenase (S6PDH) (Kim et al., 2015). In chloroplast cells, sorbitol is also hydrolyzed into hexose by sorbitol dehydrogenase (SDH). However, SDH and sorbitol oxidase (SOX) can convert sorbitol to fructose and glucose after sucrose is transported into the parenchymal cells (Nosarzewski et al., 2007; Yamaguchi et al., 1994). The sugar metabolism of Rosaceae plants is shown in Fig. 1 (Teo et al., 2006; Wang et al., 2009).

**Fig. 1.**
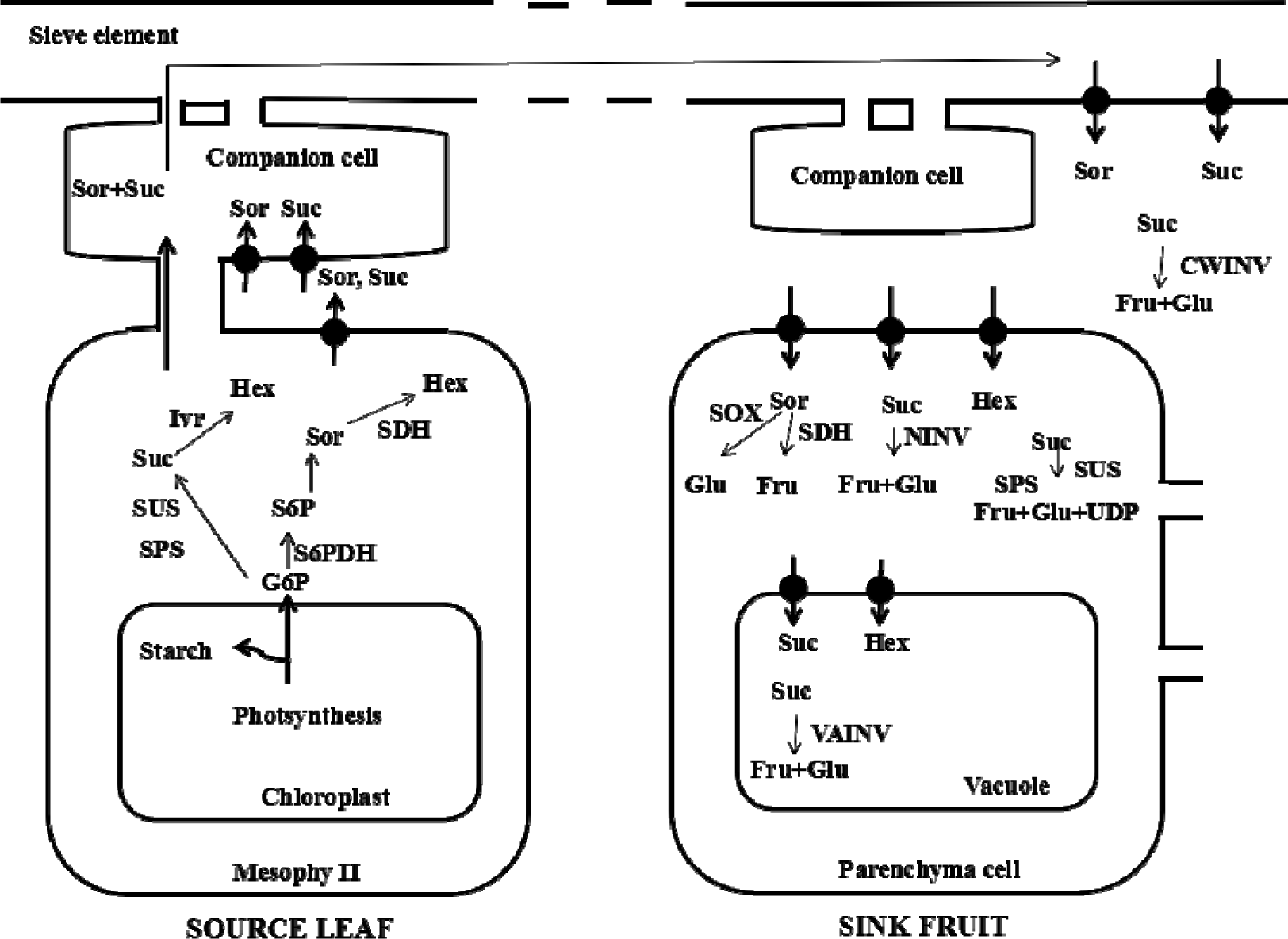
Sugar metabolic pathway of rosaceous plants. G6P, glucose-6-phosphate; S6PDH, glucitol-6-phosphate dehydrogenase; S6P, sucrose-6-phosphate; Sor, sorbitol; SUS, sucrose synthase; SPS, sucrose phosphate synthase; Suc, sucrose; Fru, fructose; Glu, glucose; Hex, hexose; SDH, sorbitol dehydrogenase; NINV, neutral invertase; VAINV, vacuolar acid invertase; CWINV, cell wall invertase; INV, invertase.

UV-B regulates plant growth, development, photosynthesis, the antioxidant system and endogenous hormones, and it affects the quantity of plants produced (Kakani et al., 2003; Wijewardana et al., 2016; Vanhaelewyn et al., 2016). Research on the postharvest quality of peaches treated with UV-B has been reported (Scattino et al., 2014). However, if the greenhouse lacks light, then the fruit quality declines. Although UV-B radiation acts as an environmental stimulus, in high doses it has detrimental effects on plant metabolism (Trentin et al., 2015). In our experiments peaches were treated with UV-B radiation during the fruiting period in 2015, and the results showed that supplementation with 1.44 kJ·m^−2^·d^−1^ of UV-B radiation can significantly improve the sugar content of peach fruit grown in a greenhouse. Based on this, little information is available in the literature on the effects of UV-B radiation on the sugar metabolism-related enzymes in a new peach variety Lumi 1. Knowledge of the control of sugar metabolism is essential to enhance fruit quality and promote fruit consumption (Desnoues et al., 2016). We investigated the changes in sugar accumulation and sugar metabolism-related enzymes when 1.44 kJ·m^−2^ d^−1^ UV-B intensity irradiated of peach trees, Lumi 1. There has been limited research on the effects of UV-B radiation on sugar metabolism in peach, thus our research has laid a foundation for future research.

## Materials and methods

### Plant material

The five-year-old peach trees (*P. persica* L. cv. Tainongtianmi) and four-year-old peach trees (*P. persica* L. cv. Lumi 1) were harvested from the same experimental and professional plantations located in an automated polycarbonate-covered greenhouse at the horticultural science experimental station of shandong agriculture university, located in Tai’an, China (117°06’ E, 36°15’ N).

### UV-B treatment

Screening for the optimum UV-B radiation dose. The using UV-B lamp tubes (30W, 297nm; Nanjing Huaqiang Electronics Co., Ltd., China), the UV-B irradiation was carried out by treatments of 0.72 kJ·m^−^·^2^d^−1^, 1.44 kJ·m^−2^·d^−1^ and 2.16 kJ·m^−2^·d^−1^ from February 2015 to May 2015. The control supplemental UV-B dose was 0 kJ·m^−2^·d^−1^. The UV-B lamp tube were hang at 0.9 m from the top of the plant. UV-B radiation doses were controlled using an electronic automatic device, from flowering to fruit maturity one hour every day from 10 a.m. to 11:00 a.m.. This program will be suspended when it was cloudy, raining or snowing. The water-fertilizer conditions of treatment group and control group were the same.

The changes in sugar accumulation and sugar metabolism-related enzymes in ‘Lumi 1’ were investigated under UV-B radiation. UV-B radiation treatments were carried out at 1.44 kJ·m^−2^·d^−1^ from February 2016 to May 2016 and supplemental UV-B radiation dose was 1.44 kJ·m^−2^·d^−1^ with supplemental 0 kJ· m^−2^·d^−1^ UV-B dose as control, and all other conditions being the same with screening for the optimum UV-B radiation dose.

Peach fruits and leaves around fruit were sampled every 14 days from the first fruit-expansion stage to the maturation stage, as Table 1 shown. The sarcocarp, pericarp and leaf samples were immediately frozen in liquid nitrogen and then stored at −75°C for later use.

**Table 1.** Sample collection times

| Sampling time | Days post-anthesis/d | Mature period |
| --- | --- | --- |
| 11-Mar | 21 | First fruit-expansion stage |
| 25-Mar | 35 | early of slow-growth stage |
| 8-Apr | 49 | late of slow-growth stage stage |
| 22-Apr | 63 | Second fruit-expansion stage |
| 6-May | 77 | Maturity stage |

### Fresh and dry weight measurements

Ten peach fruits were randomly sampled and weighed, then kiln dried, and the water content was calculated.

### Total anthocyanin extraction and measurement

Anthocyanins were extracted from fresh peach sarcocarp and pericarp in 95% ethanol (pH2) and measured spectrophotometrically at 530 nm, 620 nm and 650 nm. The anthocyanin content was calculated using the following equations:

D_λ_ = (A530-A620) - 0.1(A650-A620) and
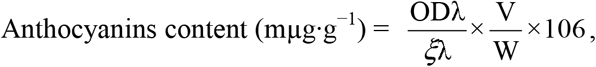

where ξλ (Anthocyanin molecule’s 530 nm light extinction coefficient) = 4.62 ×104, V indicates the total volume of the extract; W indicates the sample weight (g FW: Fresh weight); and 106 is the number of molecules of mg in mμg.

### Chlorophyll and carotenoid extractions and measurements

Chlorophyll and carotenoid were extracted from fresh pulp, peel and leaf in 96% ethanol and measured spectrophotometrically at 665 nm, 649 nm and 470 nm. The chlorophyll and carotenoid contents were calculated using the following equations:

Ca = 13.95 D665 - 6.88 D649;

Cb = 24.96 D649 - 7.32 D665;

1000 D470- 2.05 Ca - 114.8 Cb

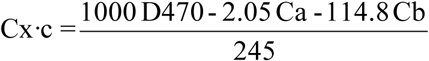; and

Pigment content 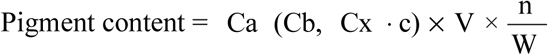
where Ca represents the concentration of chlorophylla; Cb represents the concentration of chlorophyllb; Cx·c represents the concentration of carotenoid; V indicates total volume of extract; W indicates the sample weight (g FW); and n indicates the dilution factor.

### Total phenolics and phenolic components extractions and measurements

Peach sarcocarp and pericarp were used in extractions under the following conditions previously selected (Liu et al., 2015) with slight modifications. Samples were extracted in acidic methanol-water (50:50, v/v, pH 2). Then, they were extracted in 70% aqueous acetone. Phenolic compounds that were extracted were stored at −20°C until being used in the next 24 h. The Folin–Ciocalteu method was used to determine the total phenolic content of the extracts (Singleton and Rossi, 1964). Each treatment consisted of three biological replicates, and each sample was measured using three technical replicates. The data from a typical experiment are presented.

High-performance liquid chromatography (HPLC) was used for the identification and quantification of mature phenolic components of peach sarcocarp and pericarp in a Waters series 600 chromatography unit (HPLC; Waters, Milford, MA, USA) equipped with two 515 pumps and a 2487 dual UV detector (Waters Alliance 2695 HPLC) at a wavelength of 280 nm. The chromatographic separation of phenolic components was carried out using eluent A, acetic acid aqueous solution (pH 2.8), and eluent B, acetonitrile, as the mobile phase, with a flow rate of 1.0 mL per min. Samples were injected onto a Kromasil C18 (250 × 4.6 mm) column, and the gradient was programmed as follows: 90–65% B (0–30 min), 65% B (30–40 min), 65–90% B (40–42 min) and 90% B (42–45 min). Operating conditions were as follows: column temperature was 35°C; injection volume was 10 μL, and the sensitivity was 0.5 aufs. Each treatment consisted of three biological replicates, and each sample was measured using two technical replicates. Data from a typical experiment are presented.

### Sugar and organic acid compound extractions and measurements

Samples were extracted from fresh pulp, peel and leaf in 85% ethanol and then dried. Dry solids were reconstituted in 2 mL of deionized water, and measured by HPLC for identification (Liu et al., 2015).

The chromatographic separation of sugars of pulp, peel and leaf were implemented using acetonitrile–water (75:25, v/v) as the mobile phase with a flow rate of 0.8 mL per min and a 5.0 μm NH_2_ (4.6 mm × 250 mm) column (GL Sciences Inc., Torrance, CA, USA). The samples’ chromatographic separations were performed on an YMC-Pack Polyamine II (4.6 mm × 250 mm) column. Eluted peaks were detected with an RID-10A refractive index detector (Shimadzu Co., Kyoto, Japan). Operating conditions were as follows: column temperature was 30°C, injection volume was 10 μL and the sensitivity was 4 aufs.

Organic acid analyses of pulp were carried using a Waters series 515 chromatography unit (HPLC) equipped with two 515 pumps and a 2487 dual UV detector at a wavelength of 210 nm. The chromatographic separation of organic acids was carried out using NH4H2PO4 (10 mmol/L, pH 2.3) and methanol (98:2, v/v) as the mobile phase, with a flow rate of 0.8 mL per min, and samples were injected onto a Thermo Hypersil GOLD aQ (4.6 mm × 250 mm) column. Operating conditions were as follows: column temperature was 28°C, injection volume was 10 μL, and the sensitivity was 0.5 aufs.

### Sample extraction and enzyme-linked immunosorbent assay (ELISA) measurements

Samples of peach sarcocarp and leaves were extracted in phosphate buffered saline (pH 7.4; TransGen Biotech, Beijing, China) (1:9, m/v) and fully homogenized. Enzyme liquid was stored at −20°C for no more than 7 days before being used.

Sugar metabolism-related enzyme activities, including SUS, SPS, CWINV, NINV and VAINV, and S6PDH, SDH, SOX and SOT of sorbitol metabolism-related enzyme activities, were quantified by a Plant ELISA Kit (Dong Song Bo Industry Biotechnology Co., LTD, Beijing, China), following the manufacturer’s protocol. Additionally, the same methods were used to measure the sugar transporter-related protein contents, including SUT, STP and TMT.

Pectinase activity was determined by ELISA method sames as sugar metabolism-related enzyme.

### Gene identification and primer design

The sugar metabolism-related genes investigated, *PpSUS, PpSPS, PpCWINV, PpNINV* and *PpVAINV*, and the sugar transporter genes investigated, *PpSUT, PpSTP* and *PpTMT*, were identified in Vimolmangkang et al. (2015). The sorbitol metabolism-related genes investigated, *PpS6PDH* and *PpSDH*, were identified in the NCBI database. We searched the peach genome’s annotated database to identify sugar metabolism-related genes (Verde et al., 2013). The coding sequences of these genes were further compared against the peach draft genome using the BlastN algorithm, and the genome sequences of sugar metabolism-related genes in peach were downloaded from P. persica v2.1 of the Phytozome database (https://phytozome.Jgi.Doe.gov/pz/portal.html). Primer 3 (http://bioinfo.Ut.ee/primer3–0.4.0/) was used for primer design. Primer sequences for this study are shown in Table S1.

### RNA isolation and qRT-PCR analyses

Total RNA was extracted from 500 mg of sarcocarp, pericarp and leaves using an RNAprep Pure Plant Kit (Polysaccharides & Polyphenolics-rich; Tiangen, Beijing, China). The single-stranded cDNAs were synthesized from 1 μL of RNA using a Prime Script RT reagent kit with gDNA Eraser (Takara, Dalian, China). The qRT-PCR was performed with a gene-specific primer pair, and the peach gene *GADPH* as an internal control (Tong et al., 2009). Reactions were performed on a CFX96 real-time PCR detection system with SYBR Premix Ex Taq (Takara) following the manufacturer’s instructions. The thermo-cycling parameters were as follows: 30 s at 95°C, followed by 40 cycles of 10 s at 95°C for denaturation and 40 s at 60°C for annealing and extension (Vimolmangkang et al., 2015). The specificity of the PCR was assessed by the presence of a single peak in the dissociation curve after the amplification and by the size estimation of the amplified product. The comparative cycle threshold (CT) method (2^−ΔΔCT^) method was used to quantify cDNAs with amplification efficiencies equivalent to that of the reference actin gene (Graeber et al., 2011). Each experiment was repeated at least three times using the same cDNA source.

The fluorescence quantitative CT values of *PpCWINV1,2,3,6, PpNINV5,6* and *PpTMT3,4* from leaf, pericarp and fruit were greater than 35; therefore, we assumed that they were not expressed in the tested peach variety.

### Statistical analyses

All assays were conducted with three or more biological replicates. The statistical analysis was performed using SPSS for Windows version 19 (SPSS) (Chicago, IL, USA). Categorical variables were expressed as frequencies, and percentages and continuous variables were expressed as means ± SEs, as appropriate. The data were analyzed by an analysis of variance and, when appropriate, Duncan’s test was used. A significance level of P < 0.05 was applied.

## Results

### 1.44 kJ·m^−2^·d^−1^ UV-B radiation increased the sugar content and decreased the acid content

1.44 kJ·m^−2.^d^−1^ UV-B radiation was the most effective in increasing the sugar content and reducing the acid content of the peach variety ‘Tainongtiamnmi’ in the three UV-B radiation intensities used in this study (Fig. 2). Its total sugar content was 1.5 times as much as that of the control.

**Fig. 2.**
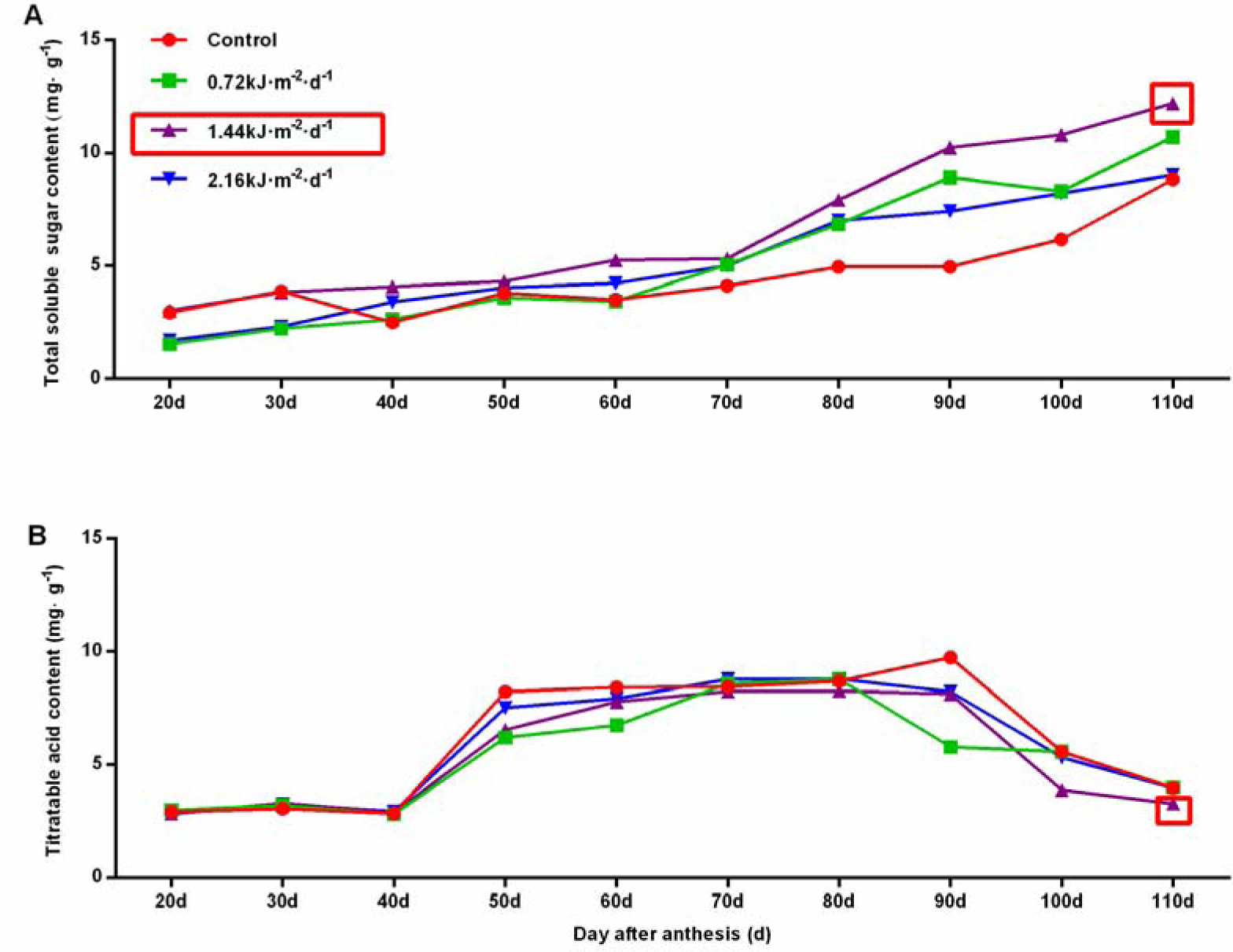
Effects of different UV-B radiation intensities on the total sugar content in peach (cv. Tainongtianmi) fruit. (A) Changes in the total sugar content after three different intensities of UV-B radiation. (B) Changes in the titratable acid content after three different intensities of UV-B radiation.

### Quality and appearance of ‘Lumi 1 ’during the UV-B treatment

The 1.44 kJ·m^−2.^d^−1^ UV-B radiation did not produce obvious effects on fresh weight, dry weight or the water content of the peach fruit (Fig. S1). Pectinase can reflect fruit softening (Santiago-Doménech et al., 2008). Pectinase activity was first increased by UV-B treatment, then decreased activity in the mature period compared with the control (Fig. S2).

The fruit color of sarcocarp and pericarp were significantly enhanced after UV-B treatment during the ripening stage (Fig. 3A, 3B). The anthocyanin contents in flesh and peel were significantly elevated compared with those in the control (Fig. 3C, 3D). After treatment with UV-B, there was no obvious law of change in the chlorophyll content, which was mainly increased in the flesh, but decreased in the pericarp and leaf (Fig. S3). The levels of carotenoids in the flesh, peel and leaf were mainly decreased by UV-B (Fig. S3).

**Fig. 3.**
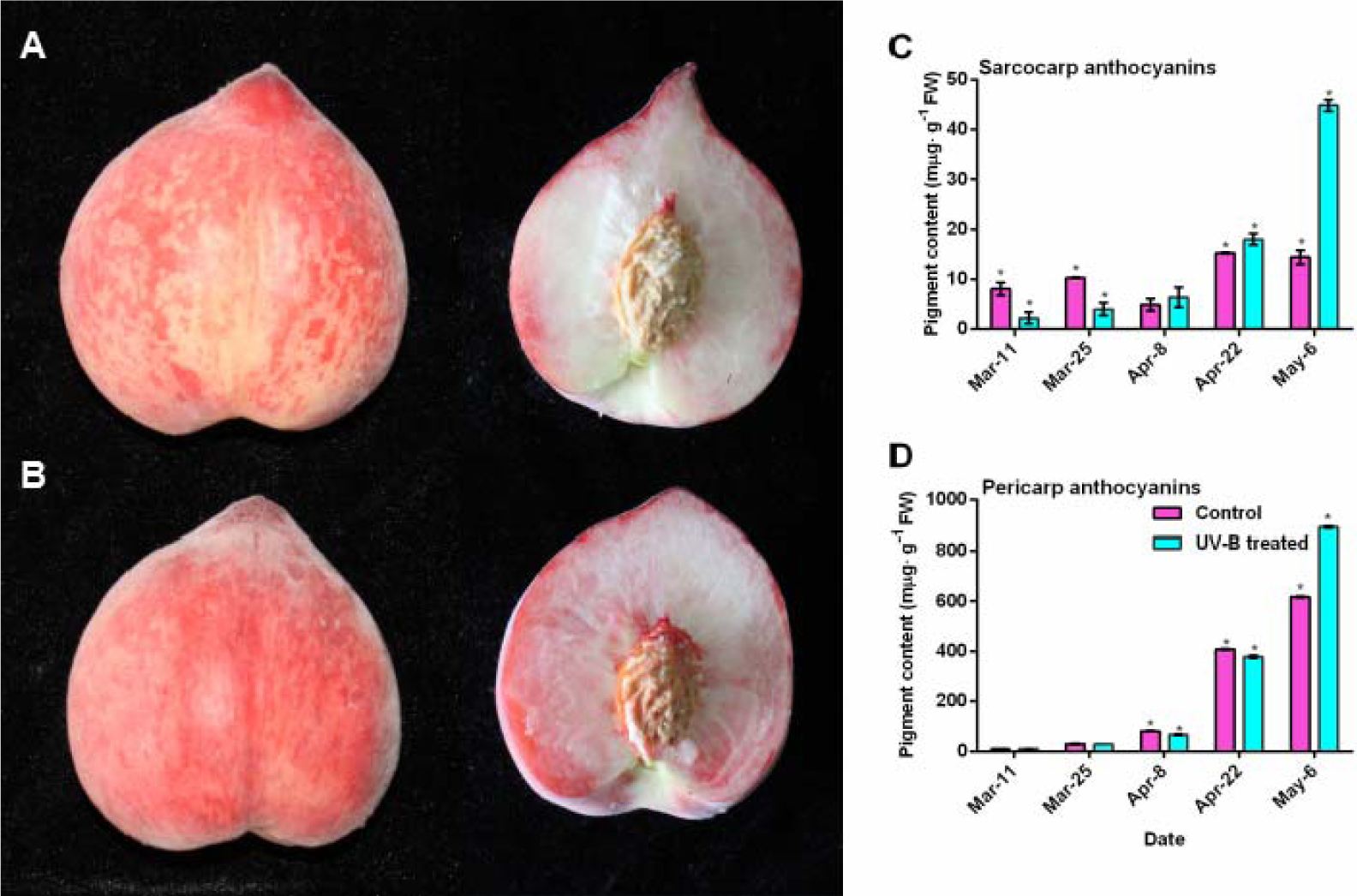
Effects of UV-B treatment on the anthocyanin contents of flesh and peel, and the mature fruit phenotype. (A) Peach fruit phenotype of control. (B) Peach fruit phenotype after UV-B treatment. (C) Changes in the anthocyanin content in fruit flesh affected by UV-B radiation (*P < 0.05). (D) Changes in the anthocyanin content in fruit peel affected by UV-B radiation (^*^P < 0.05).

Under UV-B radiation total phenolics content in the flesh was mainly increased (Fig. S4A). In the pericarp total phenolics content did not significantly change compared with in the control (Fig. S4B). At the mature stage, only the protocatechuic acid content was greater than in the control of sarcocarp, while in the pericarp, only the neo-chlorogenic acid content was greater than in the control (Fig. S4C).

### Change in sugar content during UV-B treatment of ‘Lumi 1’

After UV-B treatments, the total sucrose and sugar content levels increased compared with those of the control at the first fruit-expansion and slow-growth stages in sarcocarp (Fig. 4A). Consistently, the sugar-acid ratio was twice as much as control at late of slow-growth stage stage (Fig. 4B). Fructose and glucose in pulp were not significantly changed (Fig. 4C). The organic acid components and total organic acid content were decreased by UV-B at the slow growth phase (Fig. S5).

**Fig. 4.**
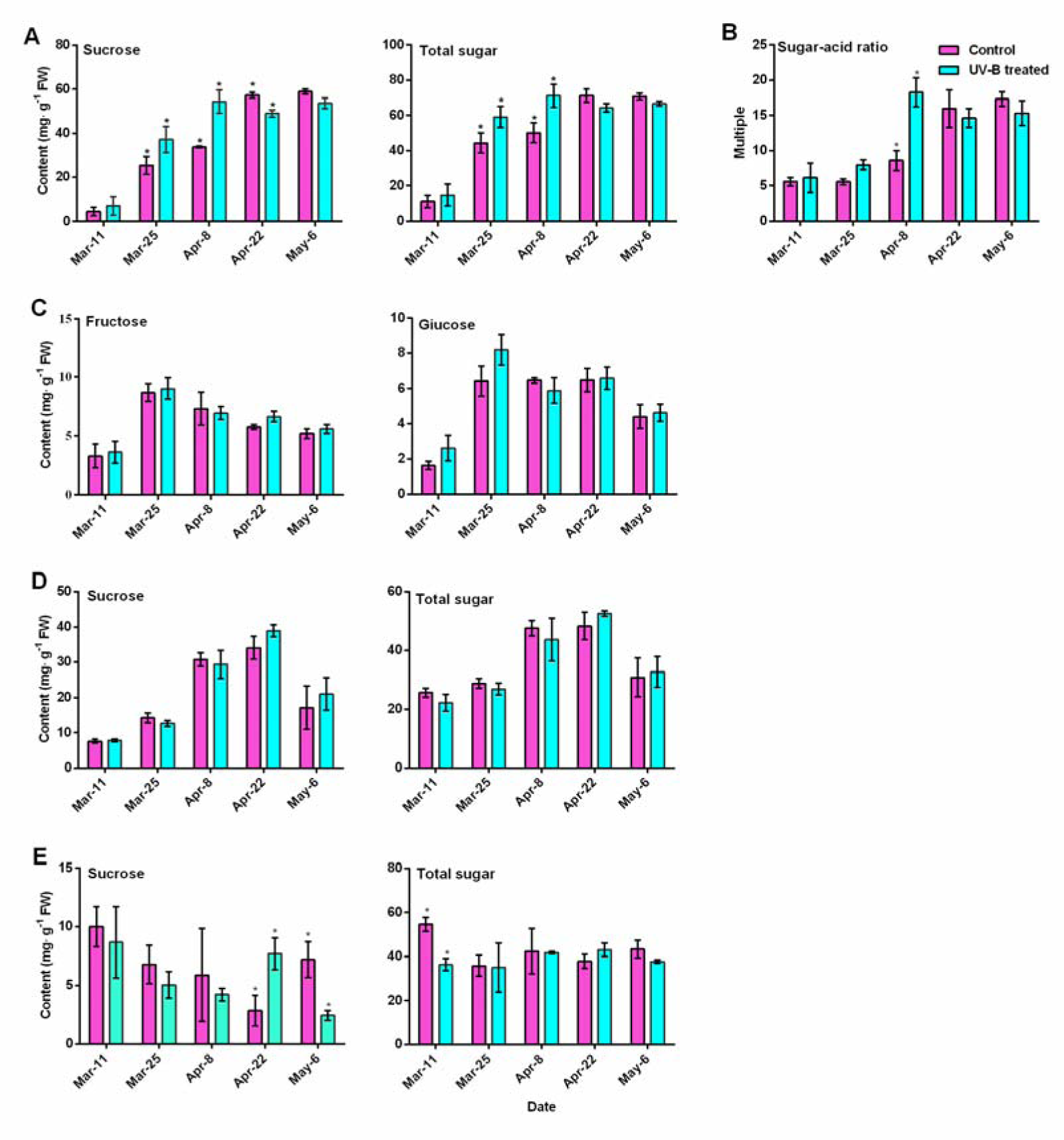
Effects of UV-B on the sugar content in different peach tissues. (A) Sucrose and total sugar contents in sarcocarp (*P < 0.05). (B) Sugar-acid ratio in sarcocarp (*P < 0.05). (C) Fructose and glucose contents in sarcocarp (*P < 0.05). (D) Sucrose and total sugar contents in pericarp (*P < 0.05). (E) Sucrose and total sugar contents in leaf (*P < 0.05).

During UV-B treatment the changes in the sucrose and total sugar contents in peel were not significant (Fig. 4D). The sucrose and total sugar contents in leaf mainly decreased, but the changes were not significant (Fig. 4E). The fructose and glucose contents in the pericarp and leaf were mainly reduced by UV-B (Fig. S6).

Thus, the effects on sucrose metabolism were divided into two stages, before the second fruit-expansion stage and after the slow growth period.

### Changes in sucrose metabolism and transport before the second fruit-expansion stage

The peach pulp is part of the sink tissue, therefore, the sucrose in the pulp cell was decomposed into fructose and glucose by SUS, SPS and INV (Fig. 1). Under UV-B treatment the SPS enzyme activity increased was up-regulated by *PpSPS2* and *PpSPS4* gene expression levels before the second fruit-expansion stage (Fig. 5A). Under regulation of four *PpSUS* genes the SUS activity was first raised and then decreased (Fig. 5B). *PpCWINV4* and *PpCWINV5* gene expression levels were decreased, but there were no significant declines in CWINV enzyme activity (Fig. 5C). Expressions of two *PpVAINV* genes were increased, consistently, VAINV activity was rised compared with the control (Fig. 5D). Expression levels of six *PpNINV* genes were down-regulated by UV-B; therefore, the NINV activity was decreased (Fig. 5E). SUT content and the *PpSUT* genes’ expression levels were decreased by UV-B compared with the control (Fig. 6A). STP’s protein content was first increased, and then decreased under the regulation of two *PpSTP* genes (Fig. 6B). TMT protein content was decreased by UV-B, by two *PpTMT* genes regulation (Fig. 6C).

**Fig. 5.**
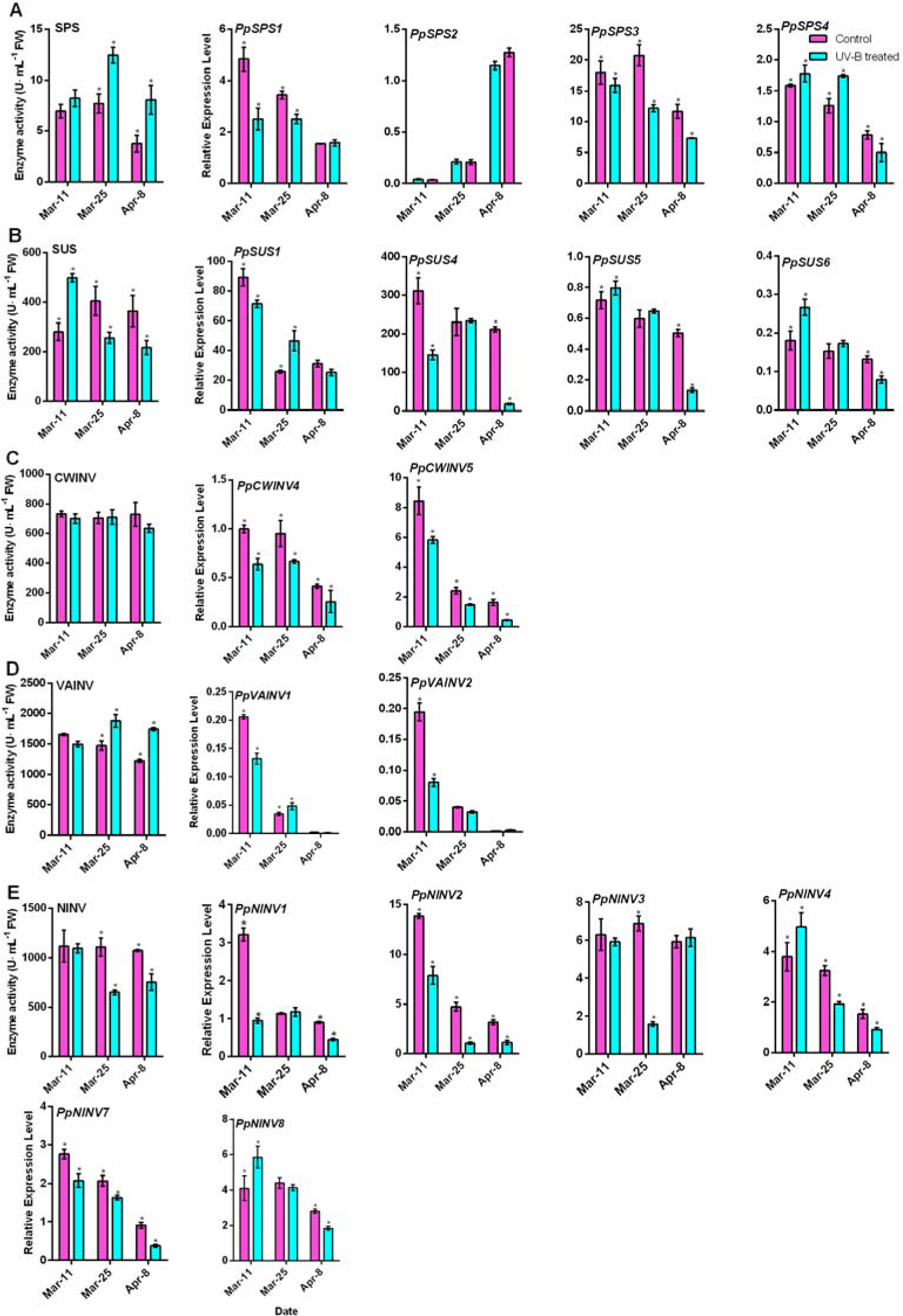
Effects of UV-B treatment on sucrose metabolism enzymes in sarcocarp before the second fruit-expansion stage. (A) SPS activity and gene expression levels (*P <0.05). (B) SUS activity and gene expression levels (*P < 0.05). (C) CWINV activity and gene expression levels (*P < 0.05). (D) VAINV activity and gene expression levels (*P < 0.05). (E) NINV activity and gene expression levels (*P < 0.05).

**Fig. 6.**
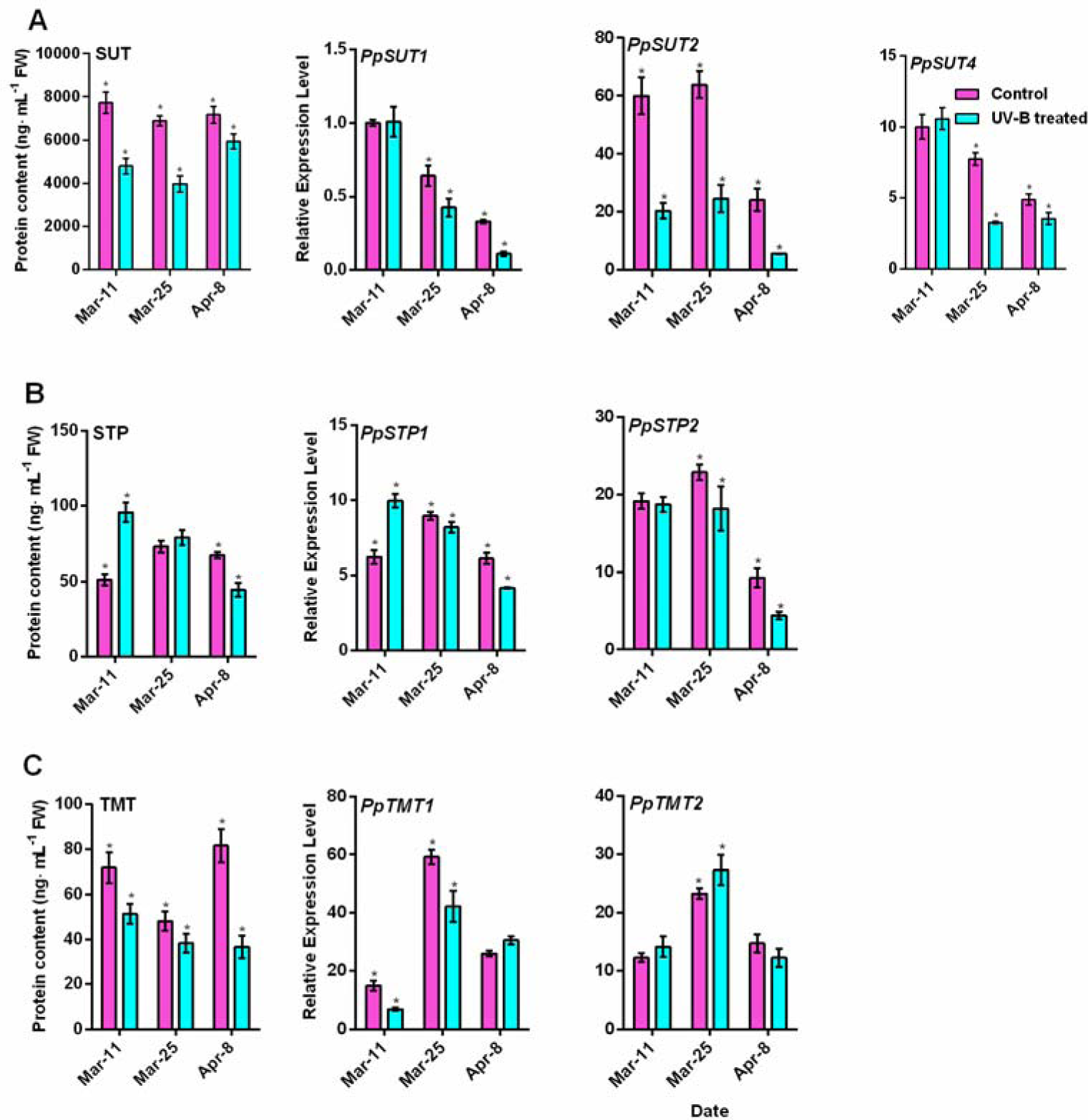
Effects of UV-B treatment on the sugar transporter before the second fruit-expansion stage in sarcocarp. (A) SUT activity and gene expression levels (*P < 0.05). (B) STP activity and gene expression levels (*P < 0.05). (C) TMT activity and gene expression levels (*P < 0.05).

SPS activity was first increased and then decreased under the regulation of four *PpSPS* genes of pericarp during UV-B treatment (Fig. 7A). SUS activity was significantly higher than in the control under the regulation of four *PpSUS* genes (Fig. 7B). Additionally, the SUT protein contents in the first two periods was significantly increased compared with the control under the regulation of three *PpSUT* genes, and then it decreased during the late of slow-growth stage period (Fig. 7C). The enzyme activities of CWINV, VAINV and NINV, the protein content levels of STP and TMT, and their gene expression levels were significantly increased by UV-B radiation (Fig. S7).

**Fig. 7.**
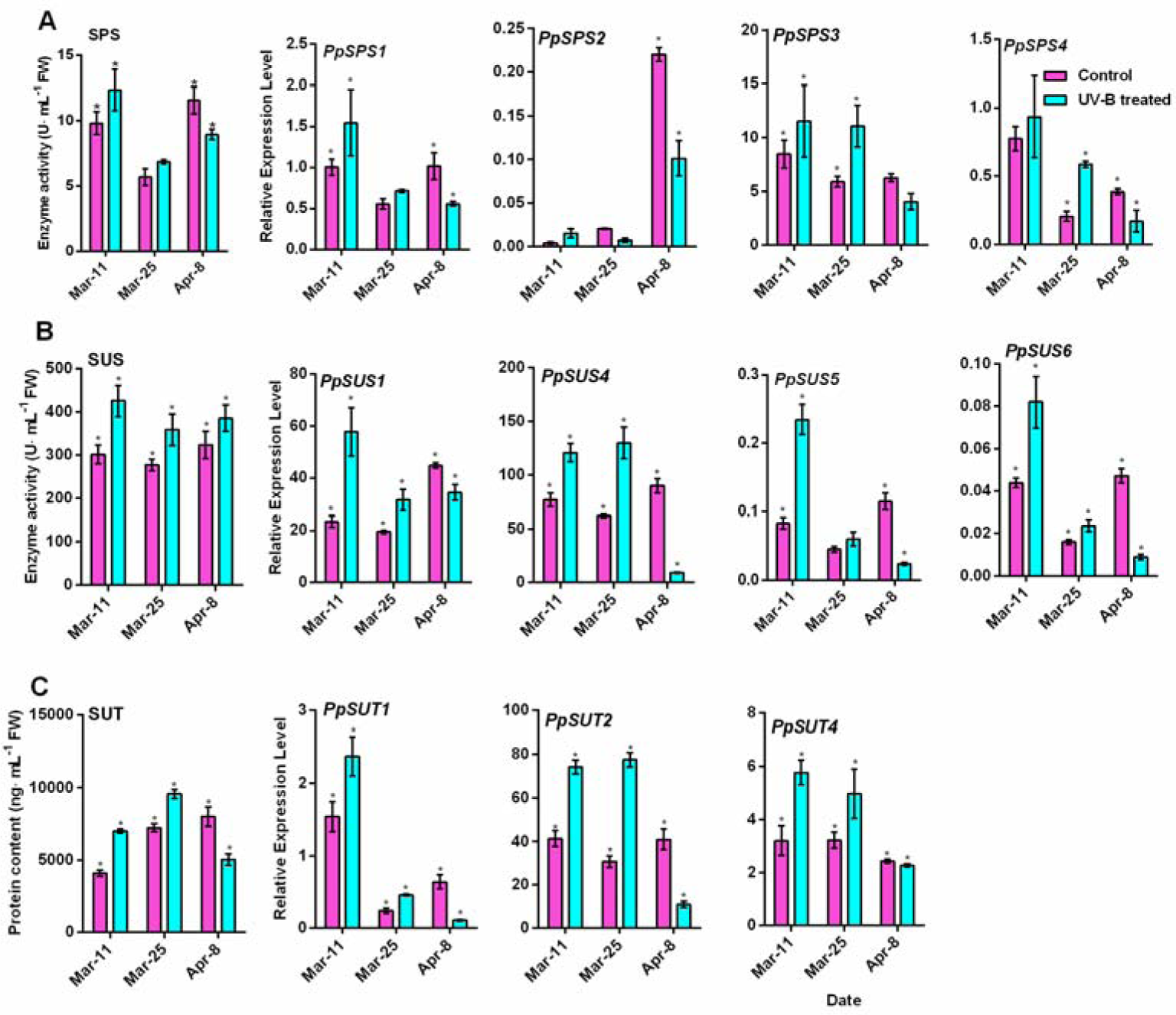
Effects of UV-B treatment on SPS, SUS and SUT activity before the second fruit-expansion stages in pericarp. (A) SPS activity and gene expression levels (*P < 0.05). (B) SUS activity and gene expression levels (*P < 0.05). (C) SUT activity and gene expression levels (*P < 0.05).

SUS and NINV enzyme activities and their gene expression levels were mainly increased of leaves during the UV-B treatment, while SPS, CWINV and VAINV enzymes and their gene expression levels did not change significantly (Fig. S8). SUT protein level first increased and then decreased, which was contrary to the accumulation of sugar in the flesh (Fig. S9A). STP and TMT protein and gene expression levels were increased compared with the control (Fig. S9B,9C).

In general, the fruit sugar content increased before the second fruit-expansion stage due to the increased synthesis and transport of sucrose in pericarp and inhibition of sucrose decomposition and transport in sarcocarp. It had little correlation with the synthesis and translocation of sugars in leaf.

### Changes in sucrose metabolism and transport after the slow-growth period

In sarcocarp, SPS, SUS, CWINV and NINV activities were gradually decreased during UV-B treatment, which were regulated by *PpSPS, PpSUS, PpCWINV* and *PpNINV* expression levels, the VAINV activity was increased compared with that of the control after the slow-growth period (Fig. 8). The levels of SUT, STP and TMT proteins and their gene expression levels in pulp were decreased by UV-B (Fig. 9).

**Fig. 8.**
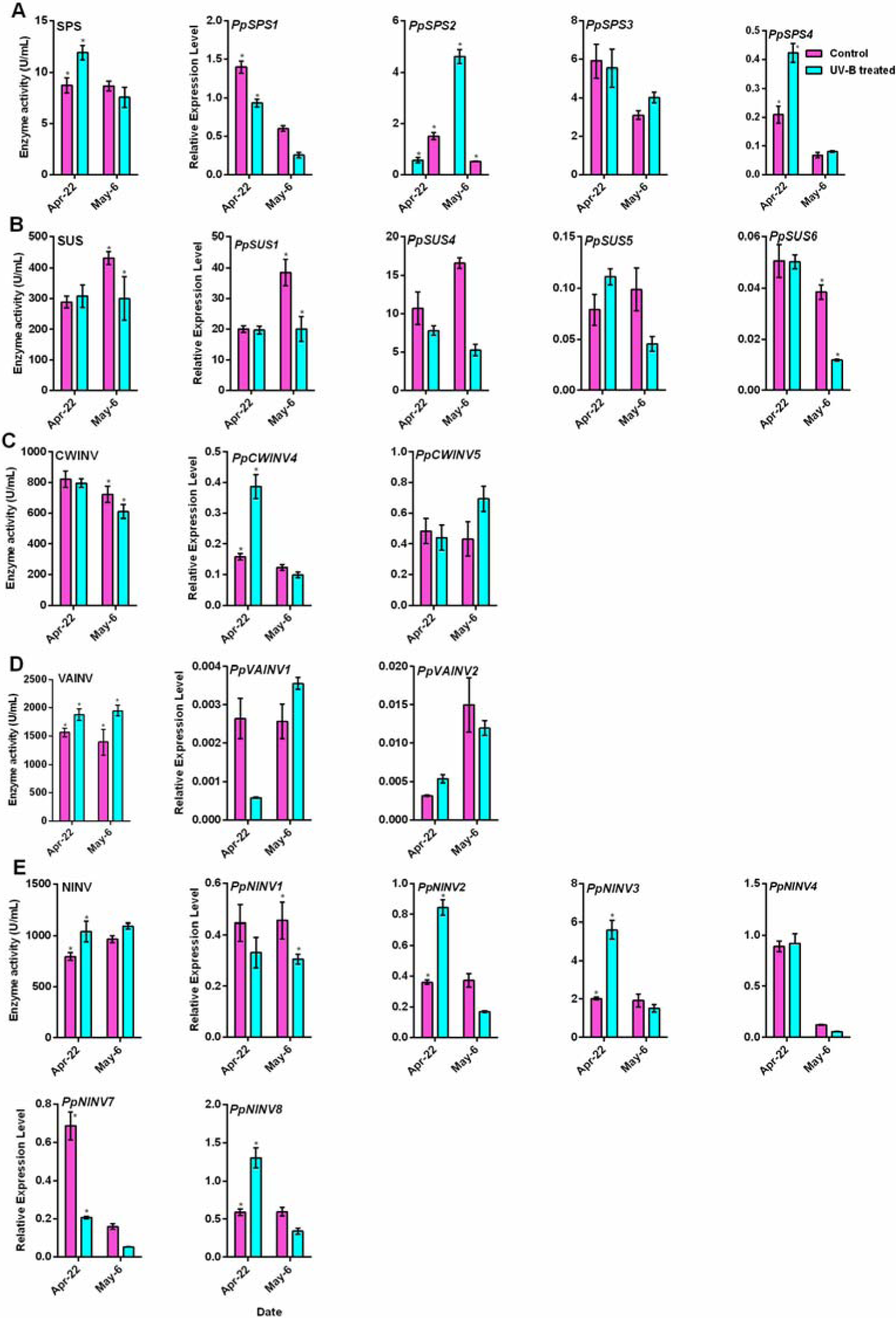
Effects of UV-B radiation on sucrose metabolism enzymes after the slow growth period in sarcocarp. (A) SPS activity and gene expression levels (*P < 0.05). (B) SUS activity and *PpSUSs* expression levels (*P < 0.05). (C) CWINV activity and gene expression levels (*P < 0.05). (D) VAINV activity and gene expression levels (*P <0.05). (E) NINV activity and gene expression levels (*P < 0.05).

**Fig. 9.**
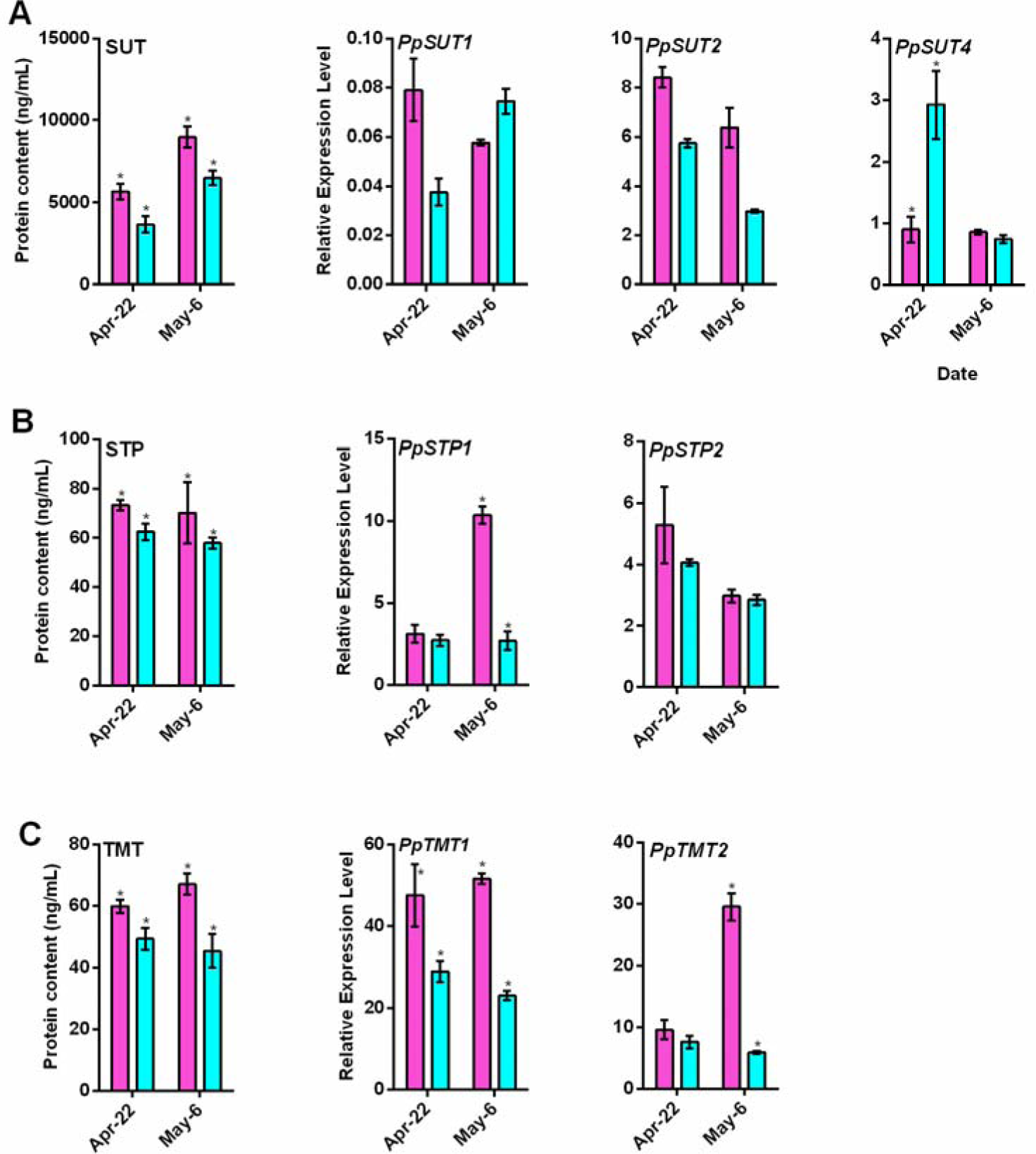
Effects of UV-B radiation on sugar transporters after the slow growth period in sarcocarp. (A) SUT activity and gene expression levels (*P < 0.05). (B) STP activity and gene expression levels (*P < 0.05). (C) TMT activity and gene expression levels (*P < 0.05).

In pericarp, the SPS and SUS activities were decreased regulated by four *PpSPS* and four *PpSUS* gene expression during UV-B treatment (Fig. 10A, 10B). SUT level and three *PpSUT* expression levels were also decreased (Fig. 10C). CWINV activity and gene expression were decreased by UV-B radiation. VAINV activity, and STP and TMT levels and their gene expression levels, were increased. Changes in NINV activity were not observed (Fig. S10).

**Fig. 10.**
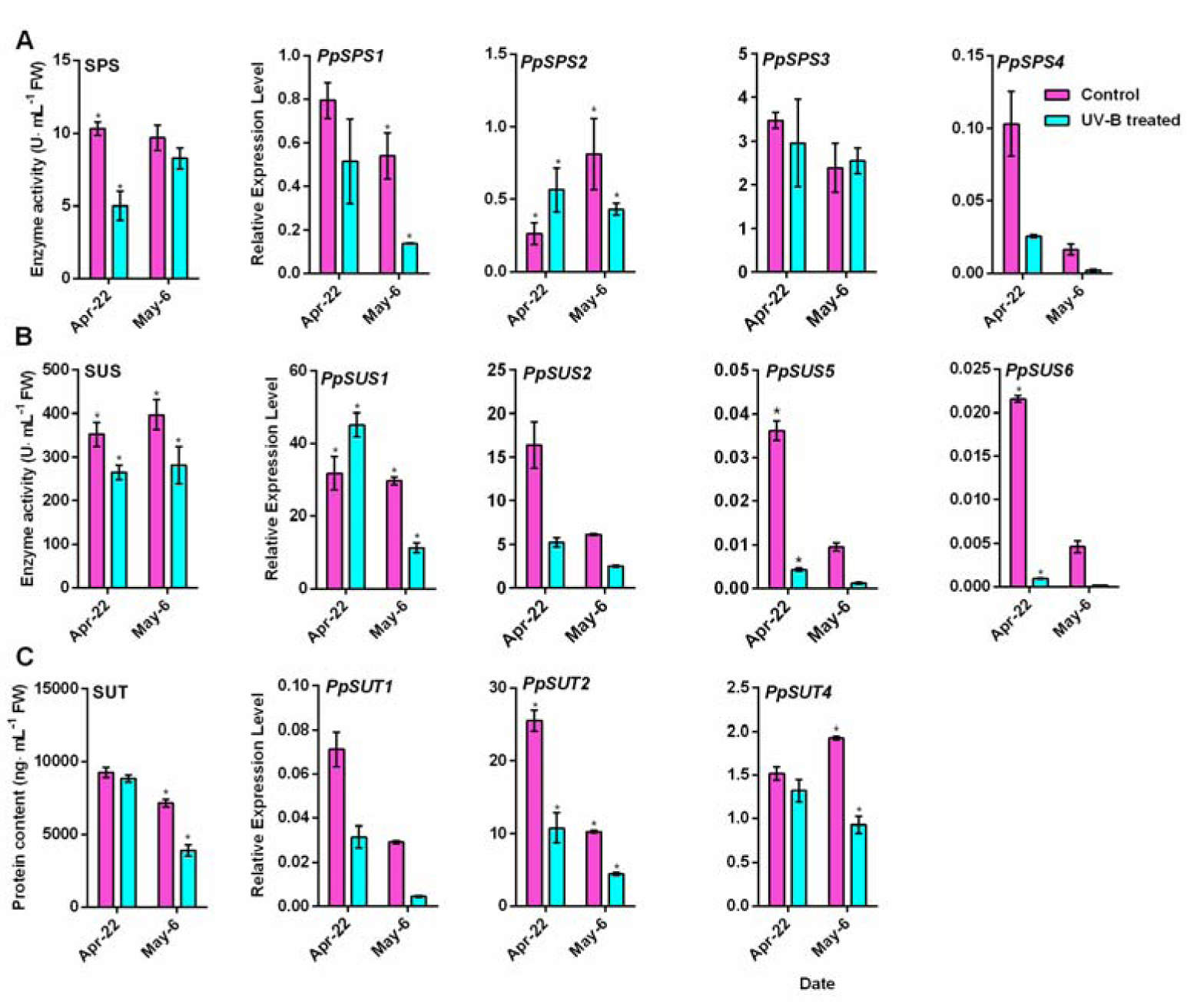
Effects of UV-B radiation on SPS, SUS and SUT after the slow-growth period in pericarp. (A) SPS activity and gene expression levels (*P < 0.05). (B) SUS activity and gene expression levels (*P < 0.05). (C) SUT activity and gene expression levels (*P < 0.05).

In leaf, regulated by *PpSPS* expression the SPS activity gradually decreased. SUS, CWINV and NINV activities and their gene expression levels were decreased by UV-B treatment. Only VAINV’s activity and *PpVAINVs* expression level were increased during UV-B treatment (Fig. S11). The transport of sugars, SUT’s protein level and the *PpSUT* expression level were decreased by UV-B radiation, while STP and TMT protein levels, and *PpSTP* and *PpTMT* gene expression levels were

On the whole, sugar decreased in fruit after the slow-growth period, which was cause by the inhibition of sucrose decomposition and transport in pericarp and leaf.

### UV-B radiation increased the sorbitol content of fruit

During the UV-B treatment, in the late of slow-growth stage and mature stages, the sorbitol content in fruit was significantly higher than in the control (Fig. 11 A). Compared with the control, sorbitol content in the peel and leaf was first decreased and then increased, the changes were not significant (Fig. 11B, 11C).

**Fig. 11.**
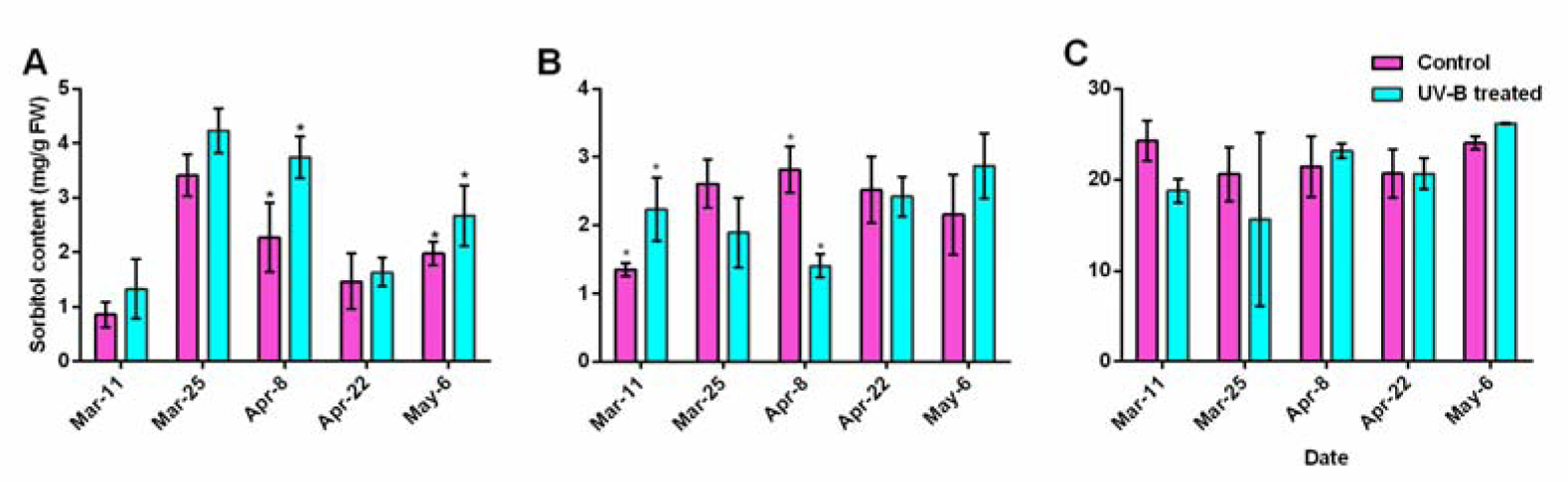
Effects of UV-B radiation on the sorbitol contents. (A) Changes in the sorbitol content of sarcocarp (*P < 0.05). (B) Changes in the sorbitol content of pericarp (*P < 0.05). (C) Changes in the sorbitol content of leaf (*P < 0.05).

In sarcocarp, SDH activity and *PpSDH* expression were first decreased and then increased compared with in the control (Fig. 12A). The SDH activity of pericarp was only significantly greater than the control at the early of slow-growth stage. Its enzymatic activity were decreased in the first fruit-expansion and mature stages compared with in the control. However, the enzymatic activity was increased in the other periods, under the regulation of *PpSDH* (Fig. 12B). In leaf, SDH activity was lower than in the control at the mature stage, but this did not correlate with the expression levels of *PpSDH* (Fig. 12C). The sorbitol content was inversely proportional to the SDH activity in sarcocarp, pericarp and leaf. In leaves, the SOT level was greater than in the control, facilitated by the transport of sorbitol to sink fruits (Fig. 13). S6PDH and SOX activities and the sorbitol content were not clearly regulated (Fig. S13).

**Fig. 12.**
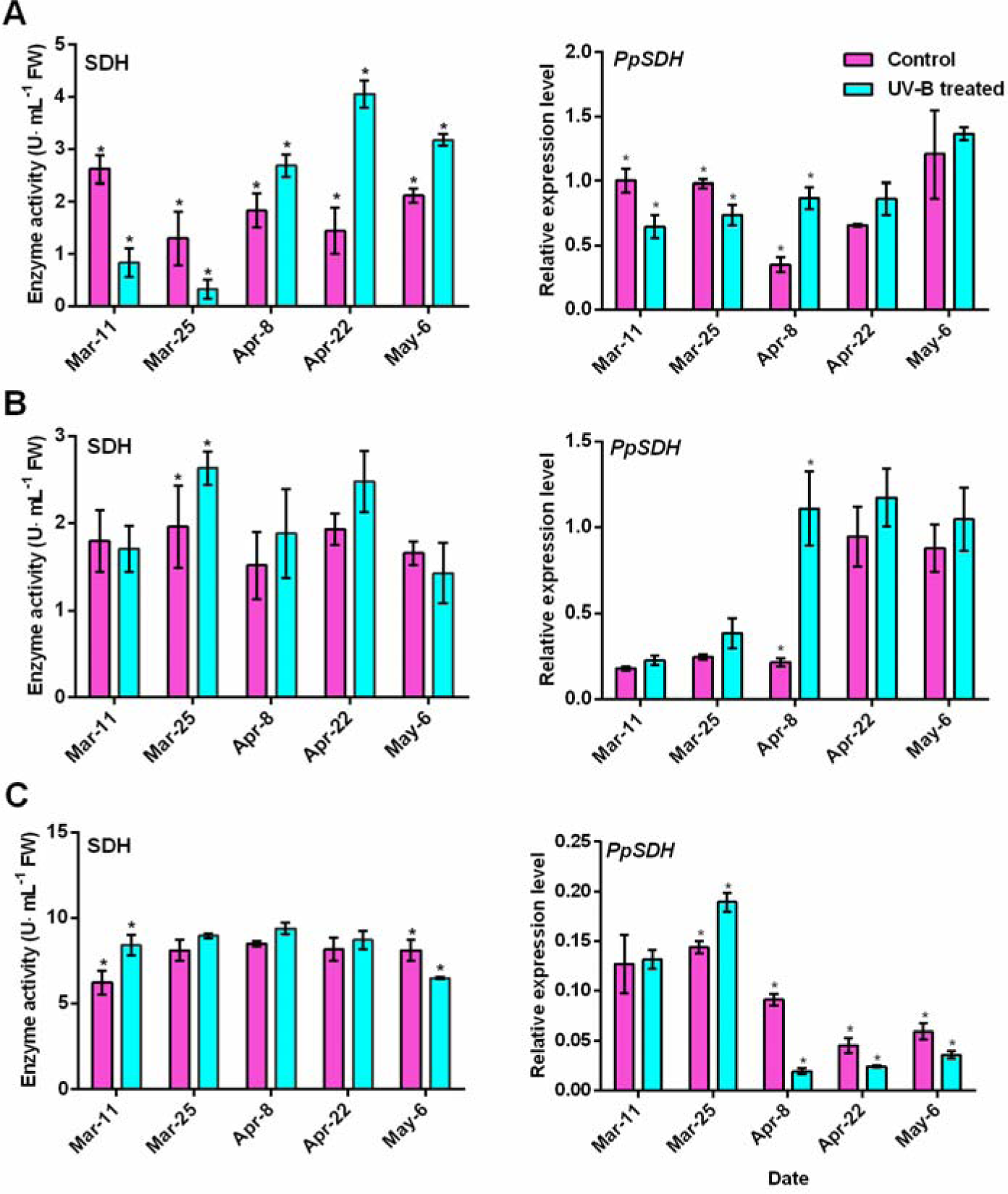
Effects of UV-B radiation on SDH activity and *PpSDH* expression levels. (A) SDH and *PpSDH* in sarcocarp (*P < 0.05). (B) SDH and *PpSDH* in pericarp (*P < 0.05). (C) SDH and *PpSDH* in leaf (*P < 0.05).

**Fig. 13.**
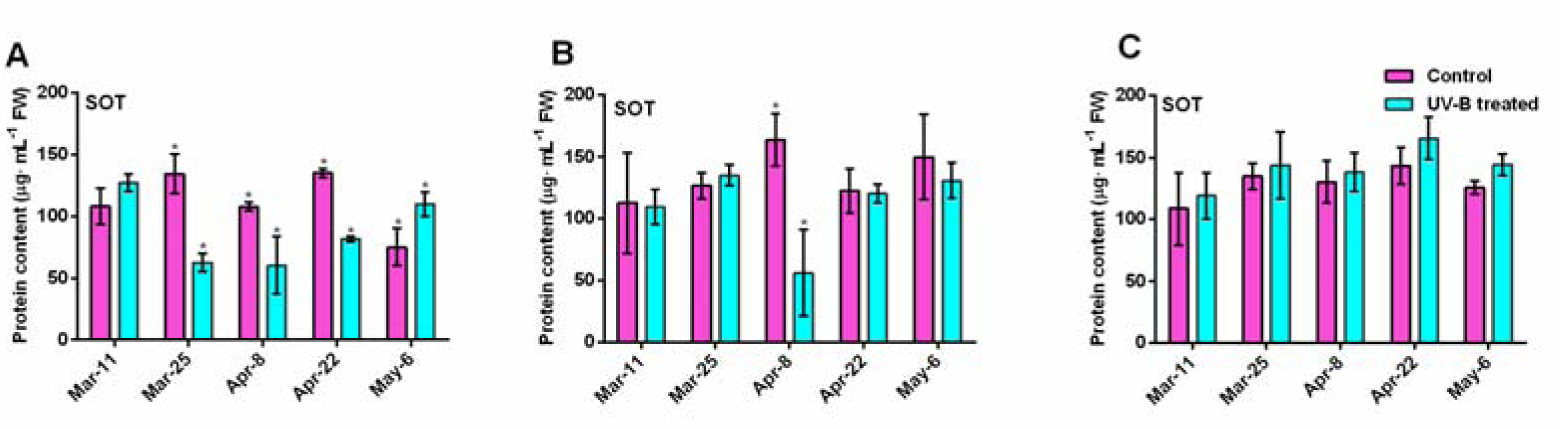
Effects of UV-B radiation on SOT activity levels. (A) SOT in sarcocarp (*P < 0.05). (B) SOT in pericarp (*P < 0.05). (C) SOTin leaf (*P < 0.05).

## Discussion

### UV-B and fruit quality

Recently, greenhouse peach consumption has been declining because of the fruit’s poor flavor, which falls short of consumer expectations (Crisosto, 2002). UV-B radiation slows grape berry development and up-regulates flavonol and anthocyanin biosynthesis (Martínez-Lüscher et al., 2014). Consistently, after UV-B treatment anthocyanin contents of ripe peach fruits’ were increased in pulp and peel, with the mature fruit coloring phenotype was deepened. However, no effects of the total phenol content were observed during UV-B treatment. In peach fruit, the tendency of sugar levels is to first increase and then decrease (Vizzotto et al., 1996), identical, trends of sugar content in sarcocarp were not altered after UV-B radiation. After UV-B radiation supplemented with nitrogen, phosphorus and potassium, the soluble sugars of pea plants are significantly enhanced (Singh et al., 2015). UV-B alleviates the uncoupling effects of elevated CO2 and increased temperature, which modulates the accumulation of sugars and upregulates the anthocyanin biosynthesis of grape berry (Martínez-Lüscher et al., 2016). Unlike the sugar results in ‘Tainongtianmi’, the sugar content after 1.44 kJ·m^−2^d^−1^ UV-B radiation was significantly enhanced, the sugar content in this experiment of ‘Lumi 1’ was slightly lower than in the control at maturity, which may have resulted from the different peach varieties used. High-intensity UV-B irradiation reduces the photosynthetic capacities of plants, resulting in crop plants becoming shorter, and having lower yields and chlorophyll contents, as well as undergoing photosystem II damage and a change in the carbon partitioning in plant organs (Schumaker et al., 1997; Feng et al., 2003; Zhang et al., 2016). The sugars may have been reduced because of long-term UV-B radiation of peach fruit at after slow-growth stage.

### UV-B, sucrose metabolism and translocation

The Rosaceae plants’ sugar metabolism is shown in Fig. 1. In the source leaf, first there is the photosynthetic synthesis of glucose-6-phosphate, then SPS and SUS catalyze the synthesis of sucrose, which is decomposed into hexose under the catalysis of the INV. Sucrose and hexose are transported to sink fruit under the action of sugar transporters. In the sink fruit, sucrose is decomposed into fructose and glucose by the catalysis of SPS, SUS and INV (Teo et al., 2006; Wang et al., 2009). UV-B treatment at gene expression and biochemical levels had different effects on these carbohydrate-metabolizing enzymes in flesh, peel and leaf. Previous studies on sugar-metabolism showed that coordinated interactions among SUS, SPS and NINV were related to sucrose metabolism and accumulation in the cytosol (Vimolmangkang et al., 2015). In rice, *PpSUT2* is involved in sucrose transport across the tonoplast from the vacuole lumen to the cytosol, playing an essential role in sugar export from the source leaves to sink organs (Eom et al., 2011). *PpSUTs* are related to sucrose metabolism and accumulation in the cytosol, participate in phloem loading in leaves and are involved in sucrose accumulation in peach fruit (Zanon et al., 2014). During UV-B treatment the increased SUT protein levels was up-regulation by *PpSUTs* of peel. The *PpSTP* gene encodes a monosaccharide transporter (Fotopoulos et al., 2003), and the heterologous expression of *PpSTP1* in yeast revealed that the encoded protein catalyzes the high-affinity uptake of glucose, galactose and mannose (Rottmann et al., 2016). TMT is well-known as a vacuolar monosaccharide importer (Wormit et al., 2006). During UV-B treatment, the decreases in SUT, STP and TMT inhibited the sucrose transport from leaves and pericarp to the fruit after after the slow-growth period after the slow-growth period. Therefor, the fruit sugar content increased before the second fruit-expansion stage due to the increased synthesis and transport of sucrose in pericarp and inhibition of sucrose decomposition and transport in sarcocarp. It had little correlation with the synthesis and translocation of sugars in leaf before second fruit-expansion stage of peach fruit development. Sugars were decreased in fruit after the slow-growth period, cause by the inhibition of sucrose decomposition and transport in pericarp and leaf. This may have been caused by sink-source relationships in fruit trees, in which, at the early stages of fruit development, the competition for nutrients is greater than in mature fruits (Lenz, 1979).

### UV-B, sorbitol metabolism and translocation

Sorbitol is mainly accumulated in source leaves in peach, and it constitutes up to 80% of the total solutes involved in osmotic adjustment (Bianco et al, 2000). Sorbitol is found mainly in leaves. In Rosaceae fruit trees, such as apple, pear and peach, leaf disc experiments showed enhanced sorbitol accumulation under salt, osmotic and low-temperature stresses, and ABA-mediated S6PDH plays a role in sorbitol biosynthesis in various stress responses (Kanayama et al., 2007). SDH exhibits the highest activity on oxidized sorbitol (Aquayo et al., 2013; Jia et al., 2015). Yeast transformed with the *MdSOTs* of apple had a significant sorbitol up-take (Watari et al., 2004). During UV-B treatment, the changes in the sorbitol content were opposite to those in the SDH activity in sarcocarp, peel and leaf. The SDH activity increased when the sorbitol content decreased, identically SDH activity decreased when the sorbitol content increased. However, SDH activity was increased and the content of sorbitol did not decrease may be due to the increased SOT protein level in the leaves, which accelerated the transport of sorbitol to the fruit during the UV-B treatment after slow-growth stage of fruit development.

### UV-B and plant stress resistance

Under salt stress, *>TMT2*> was observed to increase in abundance (Pertlobermeyer et al., 2016). Oxalic acid can significantly enhance the SUS cleavage function and SOX activity, while it also increases the sugar (glucose and fructose) content (Wang et al., 2016). Sucrose and sorbitol content were increased due to the *PpTMT* expression levels and SUS and SOX activities increased in some periods of fruit development after UV-B treatment in this experiment. Therefore, we speculate that UV-B treatment may improve the resistance of peach trees but this requires further experimental verification. Greenhouse experiments have numerous limiting factors, but in the future studies under UV-B conditions, we will add other external factors, such as CO2 or nitrogen, phosphorus and potassium, to further improve the fruit quality of peaches grown in greenhouses. Related transcription factors that promote *PpSPS, PpSUS, PpSUT* and *PpSOT* genes expression during UVB treatment of peach need to be determined (Fig. 14). We hypothesize that the application in agricultural production under greenhouse conditions of 1.44 kJ·m^−2.^d^−1^ UV-B radiation to peach before the second fruit-expansion period can increase the soluble sugar content but not the organic acids content.

**Fig. 14.**
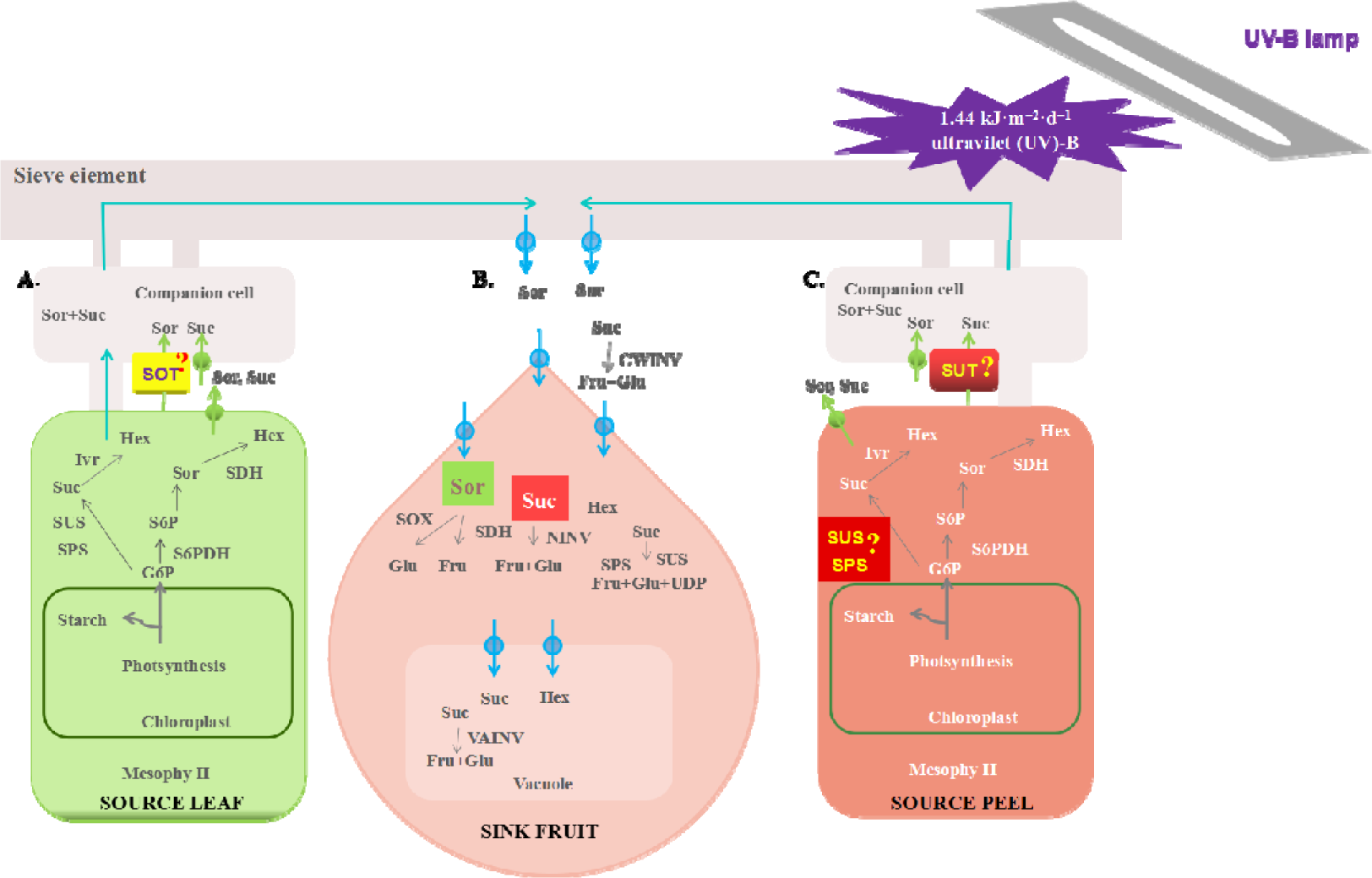
Model for the sugar metabolic pathway of peach during UV-B treatment. A. Sugar metabolic pathway of peach leves. B. Sugar metabolic pathway of peach sarcocarp. C. Sugar metabolic pathway of peach pericarp. The definition of professional term abbreviation is the same as that of Fig. 1. In the box indicates that the indicator increased during treated with UV-B. The question mark represents what we need to further determine which transcription factor is regulates the protein under UV-B treatment.

## Supplementary data

**Supplementary Table S1.** List of primers used in this study.

**Supplementary Fig. S1.** Effects of UV-B on fruit weight and water content. Supplementary Fig. S2. Effects of UV-B on the pectinase activity of sarcocarp. Supplementary Fig. S3. Effects of UV-B on chlorophyll and carotenoid contents. Supplementary Fig. S4. Effects of UV-B on phenolic acids and the total phenolic contents in the mature stage of peach development.

**Supplementary Fig. S5.** Effects of UV-B on organic acids.

**Supplementary Fig. S6.** Effects of UV-B on the fructose and glucose contents of pericarp and leaf.

**Supplementary Fig. S7.** Effects of UV-B on INV, STP and TMT before the second fruit-expansion stage in pericarp.

**Supplementary Fig. S8.** Effects of UV-B on sucrose metabolizing enzymes in leaf before the second fruit-expansion stage.

**Supplementary Fig. S9.** Effects of UV-B on sugar transporters in leaf before the second fruit-expansion stage.

**Supplementary Fig. S10.** Effects of UV-B on INV, STP and TMT after the slow-growth period in pericarp.

**Supplementary Fig. S11.** Effects of UV-B on sucrose metabolizing enzymes in leaf after the slow-growth period.

**Supplementary Fig. S12.** Effects of UV-B on sugar transporters in leaf after the slow-growth period.

**Supplementary Fig. S13.** Effects of UV-B on S6PDH and SOX changes in sarcocarp, pericarp and leaf.

## Acknowledgements

We thank Qingjie Wang for the technical assistance provided for this study. This work was supported by the National Natural Science Foundation of China (3167213) and the Natural Science Foundation of Shandong Province (ZR2014CM015).

## Author contributions

XW and XL performed the experiments and wrote the manuscript. The others provided technical support and theoretical support. LL and DG supervised the project.

